# Nanopore sequencing for the 17 modification types in 36 locations in *E. coli* ribosomal RNA enables monitoring of stress-dependent changes

**DOI:** 10.1101/2023.03.12.532289

**Authors:** Aaron M. Fleming, Songjun Xiao, Cynthia J. Burrows

## Abstract

*Escherichia coli* possess the 16S and 23S rRNA strands that have 36 chemical modification sites with 17 different structures. Direct RNA nanopore sequencing using a protein nanopore sensor and helicase brake, which is also a sensor, was applied to the rRNAs. Nanopore current levels, base calling profile, and helicase dwell times for the modifications relative to non-modified synthetic rRNA controls found signatures for nearly all modifications. Signatures for clustered modifications were determined by selective sequencing of writer knock-out *E. coli* and sequencing of synthetic RNAs utilizing some custom-synthesized nucleotide triphosphates for their preparation. The knowledge of each modification’s signature, apart from 5-methylcytidine, was used to determine how metabolic and cold-shock stress impact rRNA modifications. Metabolic stress resulted in either no change or a decrease, and one site increased in modification occupancy, while cold-shock stress led to either no change or a decrease. In the 16S rRNA, there resides an m^4^C_m_ modification at site 1402 that decreased with both stressors. Using helicase dwell time, it was determined that the *N*^*4*^ methyl group is lost during both stressors, and the 2’-OMe group remained. In the ribosome, this modification stabilizes binding to the mRNA codon at the P-site resulting in increased translational fidelity that is lost during stress. The *E. coli* genome has seven rRNA operons (*rrn*), and earlier studies aligned the nanopore reads to a single operon (*rrnA*). Here, the reads were aligned to the seven operons to identify operon-specific changes in the 11 pseudouridines. This study demonstrates that direct sequencing for >16 different RNA modifications in a strand is achievable.

## Introduction

Chemical modification of RNA occurs naturally in all phyla of life that includes over 140 different structures.^1^ These modifications are essential for RNA to adopt specific secondary and tertiary structures.^2^ They are also used for epitranscriptomic regulation,^3^ and cells employ them to mark self vs. non-self RNA and to send foreign RNA to decay pathways.^4^ The application of mass spectrometry has enabled many of these understandings, but to perform these experiments, the RNA polymers are enzymatically degraded to their nucleoside monomers for analysis.^5^ Thus, all sequencing information is lost. The sequence information for some RNA modifications is generally found one modification at a time using custom approaches to introduce specific signatures that can be found in cDNAs after high-throughput sequencing.^6,7^ Expanding this approach to inspect more than one modification at a time is challenging because it is difficult to find chemical tools that are applicable to more than one modification under similar reaction conditions.^8^ The payoff of sequencing multiple RNA modifications in a strand at one time will allow researchers to understand how these chemical decorations are used together for cellular control over phenotype and in response to stress.

Direct sequencing of RNA with a nanopore is a technology that has the potential to locate and quantify more than one modification in the strand.^9^ Nanopore sequencing of RNA works by electrophoretically passing the strand through a small aperture protein nanopore. As the nucleotides pass through the narrow zone of the protein, the ionic current is modulated based on the sequence. The ionic current levels are then deconvoluted with a recurrent neural network to yield base calls. Roughly five nucleotides of RNA contribute to the current levels analyzed, referred to as a k-mer. Success in this approach requires an ATP-dependent 3’,5’ helicase to slow the movement of the RNA through the nanopore and allow enough time to record the small current level differences. Lastly, available base-calling algorithms for RNA have been trained on canonical RNA sequences and are not modification aware. Nevertheless, chemical modifications to RNA can be identified by their unique signatures compared to the canonical forms in the base call, ionic current, and/or dwell time data as they pass the helicase active site and then through the constriction zone of the pore.^10-17^

Modifications to RNA resulting in base calling differences have enabled sequencing for pseudouridine (Ψ), *N*^6^-methyladenine (m^6^A), inosine (I), the 2′-*O*-methylation of A, C, G, and U, inosine, and N7-methylguanosine (m^7^G), as examples.^10-17^ Some RNA modifications, such as 5-methylcytidine (m^5^C) do not strongly impact the base calling and are not easily found by this method.^13^ Alternatively, the ionic current level can differ for RNA modifications, such as Ψ, as it passes through the nanopore and can be analyzed to determine the frequency of a modification.^10-12^

This approach has a challenge: RNAs do not necessarily impact the current level when centered in the k-mer, and prior knowledge of this impact is needed for quantitative analysis of the data.^12^ Another domain in which RNA modification can be revealed in nanopore sequencing is in the active site of the helicase motor protein where the translocation rate differs for the modification relative to the canonical form.^12,18^ Pseudouridine provides a recognized example of an RNA modification that impacts the helicase dwell time data.^12^ While many studies have focused on analyzing a single modification type in RNA by nanopore sequencing, there have been a few examples of multi-modification analysis. These include sequencing *E. coli* tRNA and rRNA with a nanopore to demonstrate these hypermodified RNAs produce base call data differences at known modification sites,^19^ and rRNA has been sequenced to find the 16S Ψ516 and m^7^G527 concurrently.^11^ Lastly, the four 2’-*O*-methyl ribose modifications and Ψ in rRNA have been profiled in *E. coli* and yeast.^10,18^

In the present work, we focused our sequencing efforts on *E. coli* 16S and 23S RNA because these strands have 36 well-established chemical modifications at known sites with known levels under standard growth conditions (aerobic LB media, 37 °C, grown to stationary phase).^2,20^ There exist 17 different modified RNA structures occurring on the four canonical nucleotides (A family: m^6^A, *N*^6^,*N*^6^-dimethyladenosine (m^62^A), and 2-methyladenosine (m^2^A); C family: m^5^C, 2’-*O*-methylcytidine (C_m_), *N*^4^,2’-*O*-dimethylcytidine (m^4^C_m_), and 5-hydroxycytidine (ho^5^C); G family: 2’-*O*-methylguanosine (G_m_), *N1*-methylguanosine (m^1^G), *N*^2^-methylguanosine (m^2^G), and m^7^G; U family: Ψ, N3-methylpseudouridine (m^3^Ψ), 5-methyluridine (m^5^U), N3-methyluridine (m^3^U), dihydrouridine (D or hU), and 2’-*O*-methyluridine (U_m_); Figures 1A and 1B). The goal of the present work was not to find new modifications with nanopore sequencing. Instead, it was to discover how the sequencer can be used for the inspection and quantification of a diverse set of RNA modifications on the same RNA strand. These studies are the first to catalog the base call, ionic current, and helicase dwell signatures for the 36 modifications at once. The findings are then used to address whether and how the modifications change in *E. coli* rRNA after exposure to metabolic or cold-shock stress. Lastly, the *E. coli* genome possesses 7 rRNA operons (*rrn*),^21^ allowing us to profile the operon expression levels and determine the operon-specific Ψ and m^3^Ψ modification levels before and after stress.

**Figure 1.**
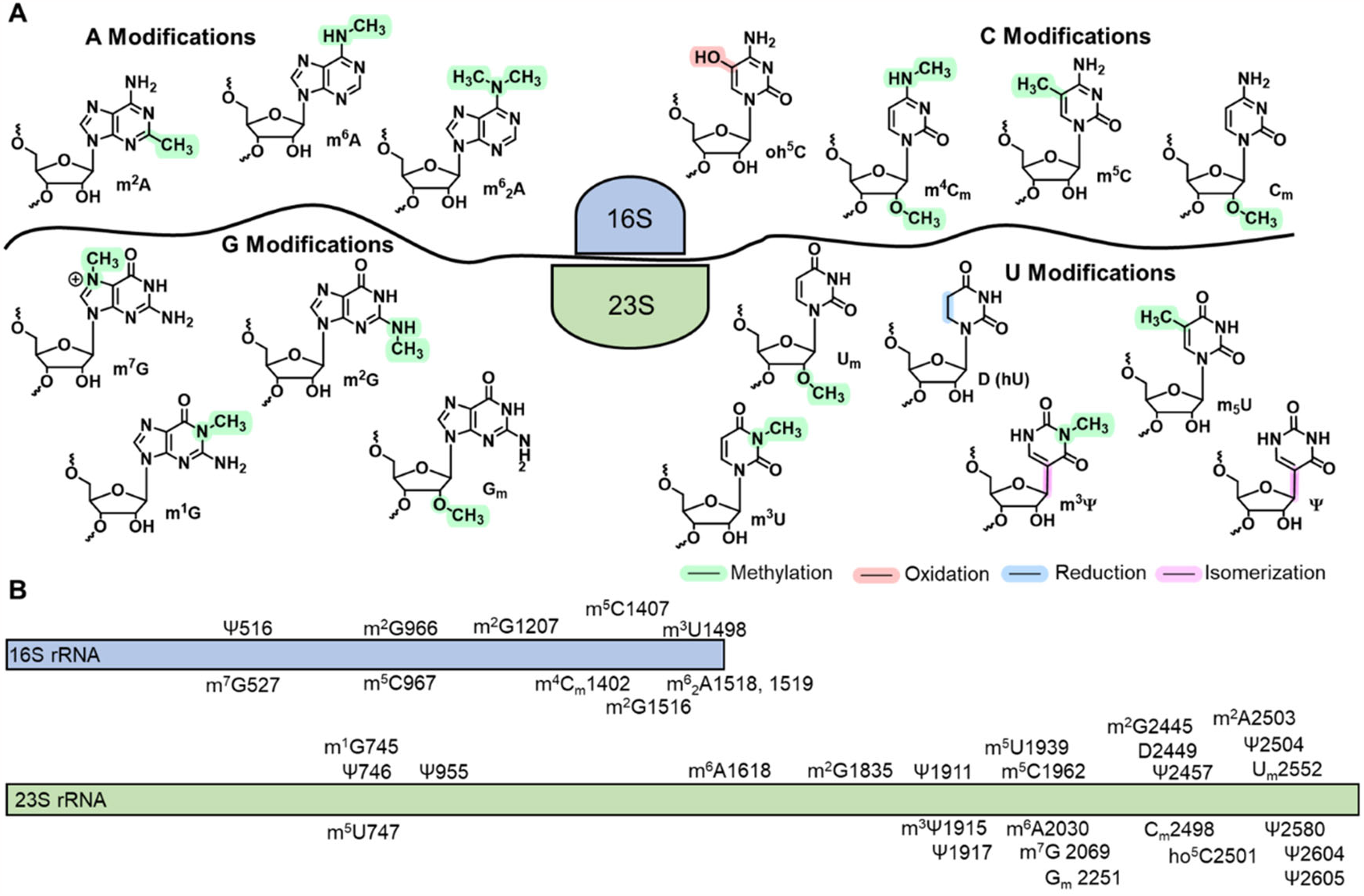
(A) Structures and (B) locations of the established RNA modifications found in the 16S and 23S rRNA strands from *E. coli*.

## Results

### Defining the sequencing signatures for the 36 RNA modifications in *E. coli* Rrna

The *E. coli* K12 DH5 alpha strain with a plasmid expressing the ampicillin resistance gene was grown aerobically in ampicillin-containing LB media at 37 °C to stationary phase after which the total RNA was harvested. The RNA integrity was verified by gel analysis, and then heat denatured for 10 min at 65 °C, flash cooled, and finally 3’-poly-A tailed using a commercial kit. The 3’-poly-A tailed RNA was the input for library preparation using the Oxford Nanopore Technologies (ONT) direct RNA sequencing kit that ligates a poly-T adaptor on the 3’ ends of the poly-A tailed RNA followed by reverse transcription to yield a DNA:RNA heteroduplex for sequencing. A final sequencing adaptor is then ligated on to the duplex. The library-prepared RNA was sequenced on either a MinION flow cell or Flongle flow cell, both with the R9.4 chemistry and reads with Q > 7 were analyzed. Synthetic 16S and 23S rRNAs with sequences from the *rrnA* operon were prepared by in vitro transcription (IVT) and sequenced to provide a control for the RNA nanopore sequencing profile without modifications. The raw sequencing data in fast5 format were base called using Guppy (6.0.7) to generate the base called data in fastq format. The reads were aligned to the 16S and 23S sequences found in the *rrnA* operon using minimap2,^22^ sorted with SAM tools,^23^ and visualized with Integrative Genomics Viewer (IGV).^24^ The base call data were quantified with Nanopore-Psu^14^ and ELIGOS2.^13^ Lastly, the ionic current and dwell time data were analyzed and quantified with Nanopolish,^25^ Nanocompore,^15^ and Tombo.^26^ All computational tools were used following their online manuals without change.

Sequencing RNA with the commercial nanopore system occurs from the 3’ to 5’ ends impacting the data in a few ways (Figure 2A).^9^ First, it is well established that there is greater read depth for the 3’-end of the sequence than the 5’-end. Second, the modifications first pass through the helicase and if they alter the translocation kinetics of the enzyme, the impact will be observed ∼11 nucleotides 3’ to the modification because this is the sequence in the nanopore sensor that will be base called and aligned to the reference sequence (Figure 2B).^12^ In the current level data, the RNA modification may or may not impact the current level when located in the center of the k-mer resulting in challenges in identifying the current level for more than one modification in proximity to one another (Figure 2C).^12^ In the base call data, the impact is usually observed at the location of the modification (Figure 2C).^13^

**Figure 2.**
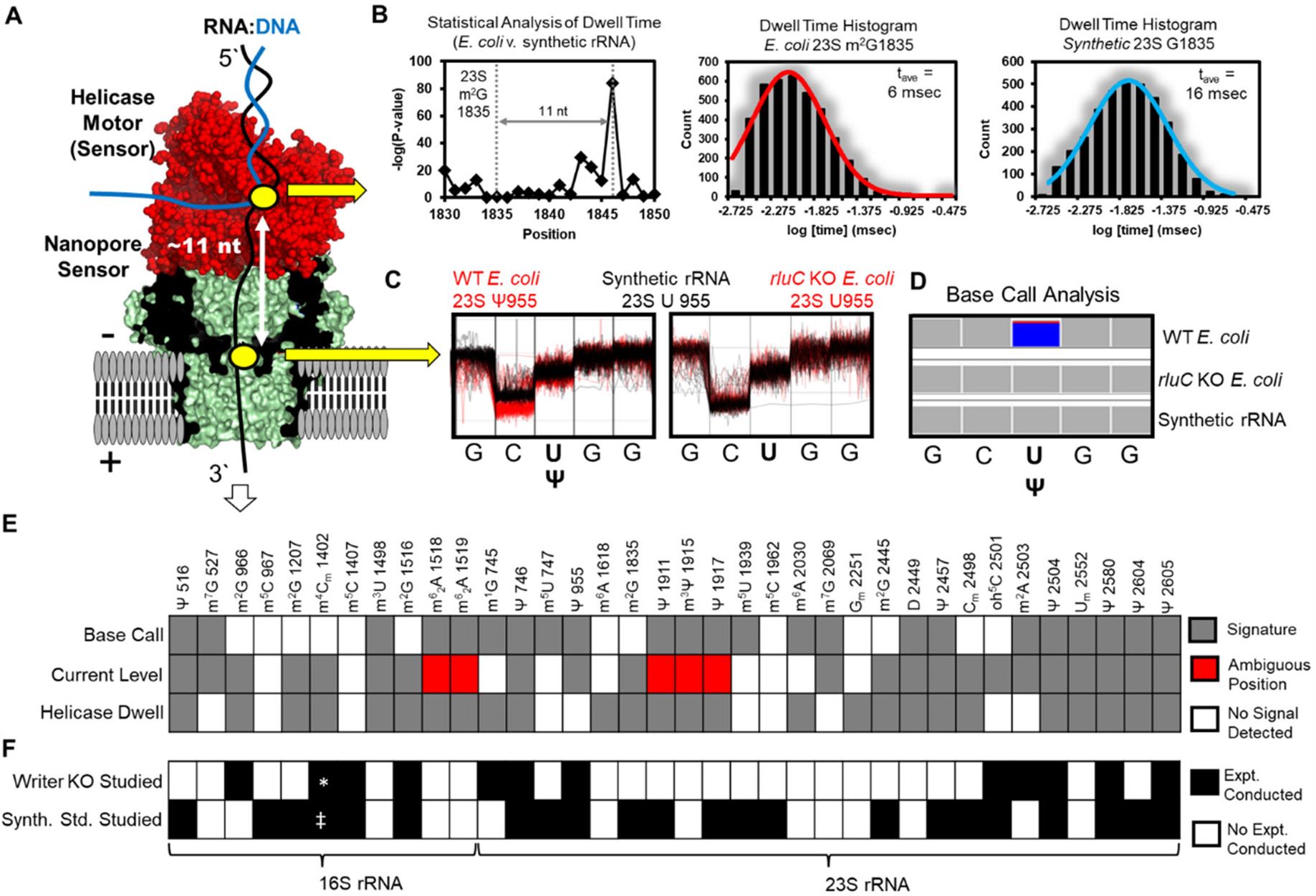
Direct RNA nanopore sequencing for chemical modifications to *E. coli* rRNA. (A) Schematic of the nanopore sequencer illustrating the protein nanopore and helicase sensor positions. (B) Helicase dwell time is altered by RNA modifications, as illustrated for 23S m^2^G1835 in *E. coli* rRNA and 23S G1835 in synthetic rRNA. (C) The nanopore ionic current levels can be impacted by RNA modifications, as shown for 23S Ψ955. This example demonstrates the current impact may not occur when centered in the 5-nt k-mer, and the signal is lost when the Ψ 955 writer protein rluC is knocked out as verification the signal comes from the modification. (D) Base calling can be impacted by an RNA modification that is illustrated in the IGV plot for the U 955 site in wild-type *E. coli* with Ψ, and the *rluC* gene knockout strain, and synthetic rRNA without Ψ at this site. In the plot, blue represents a miscall to C and gray a correct call to U relative to the reference sequence. (E) A color-coded plot of base calling error, ionic current level, and dwell time nanopore sequencing signatures for all 36 *E. coli* rRNA modifications. (F) A color-coded plot of *E. coli* knockout strains and synthetic RNAs with and without modifications that were sequenced to understand the modification signatures. *The writer for the *N*^*4*^ methyl group (rsmH) was studied for m^4^C_m_. ^‡^An RNA generated by IVT containing m^4^C was studied, and an attempt to synthesize an RNA strand by IVT with m^4^C_m_ failed.

The base call data, ionic current, and dwell time data impact observed for all 36 *E. coli* RNA modifications allowed most of them to be identified (Figure 2E). The synthetic *E. coli* rRNA was used for comparison and served as a control to eliminate any non-random errors in the data, which is an inherent feature of direct RNA nanopore sequencing data. Seventeen of the modifications are in the primary sequence at sites 5 nucleotides or more distant from other modifications, and therefore, the changes between the biological and synthetic data sets could easily be attributed to the RNA modification (16S: Ψ516, m^7^G527, m^2^G1207, m^4^C_m_1402, m^5^C1407, and m^3^U1498; 23S: Ψ955, m^6^A1618, m^2^G1835, m^5^U1939, m^5^C1962, m^6^A2030, m^7^G2069, G_m_2251, Ψ2457, U_m_2552, and Ψ2580; Figures 1, 2E gray). In a few cases, confirmation that the signals were a result of the modification came from sequencing rRNA derived from writer knock-out *E. coli* (Figures 2C and SX). Figure 2C illustrates loss of the signal from the 23S Ψ955 when its writer protein rluC is knocked out and the rRNA is sequenced. This example also shows the current level change occurs at the k-mer with the 5′ C centered in the nanopore sensor.

Nineteen of the modifications were clustered with another modification within 5 or fewer nucleotides in the sequence, and their individual signatures, particularly in the ionic current and helicase dwell time data, could not easily be determined (16S: m^2^G966 and m^5^C967; m^2^G1516, m^6^_2_A1518, and m^6^ _2_A1519; 23S: m^1^G745, Ψ746, and m^5^U747; Ψ1911, m^3^Ψ1915, and Ψ1917; m^2^G2445 and D2449; C_m_2498, ho^5^C2501, m^2^A2503, and Ψ2504; as well as, Ψ2604 and Ψ2605; Figures 1, 2E). To understand the signatures for each of these clustered modifications, RNA was sequenced from an *E. coli* strain in which an established writer for a modification was knocked out;^27-29^ the data were compared to the RNA sequencing data from the wild-type strain to allow identification of the unique signature for the members of these clustered modifications (Figure 2F). Lastly, the sequencing of synthetic RNA made by IVT with commercial m^5^CTP or CTP at seven different locations, reconfirmed that this modification does not consistently impact the sequencing data (Figure 2F).

The approach outlined enabled us to systematically determine the signatures for all modifications with a few exceptions (Figure 2E gray). In the 16S rRNA, there are two m^6^_2_A residues at positions 1518 and 1519 written into the RNA by the same methyltransferase;^28^ base call data allowed quantification, even though they gave current signatures that could not be easily deconvoluted (Figure 2E red). Second, in the 23S rRNA, there exist Ψ1911, m^3^Ψ1915, and Ψ1917 in which the isomerization of U to Ψ at the three sites is catalyzed by the same synthase, after which the methyl group on N3 of Ψ1915 is installed.^5^ It was not possible to definitively determine nanopore ionic current signatures for these modifications (Figure 2E red); nonetheless, the base call data could be used for their identification and quantification (Figure 2E gray). Finally, synthetic RNAs prepared by IVT containing m^5^C, m^4^C, C_m_, and ho^5^C were installed with their commercially available NTPs, and NTPs were synthesized for m^2^G, m^5^U, m^4^C_m_ to install them in long RNAs (Figures SX). This allowed independent verification of their base call, ionic current, and dwell time signatures in 7 different k-mer contexts (Figure 2F). Inspection of the data identified base call signatures for 24 of the modifications, nanopore ionic current level signatures for 27 of the modifications, and helicase dwell time signatures for 26 of the modifications (Figure 2E). These signatures were used in the studies that follow.

### Monitoring stress-dependent changes to rRNA modifications by nanopore sequencing

The knowledge of nanopore signatures for the rRNA modifications was used to detect and measure their occupancy in the strands under normal growth conditions and after exposure to metabolic or cold-shock stress. Prior reports have used mass spectrometry or gel analysis to quantify each of the *E. coli* rRNA modifications under stationary phase growth in LB media at 37 °C, similar to the present conditions, to find they exist at >85% occupancy and many at near quantitative levels.^20,30-33^ In the first analysis, the data from the nanopore sensor were interrogated (Figure 3A). The current levels were inspected and quantified using Tombo;^26^ recall, RNA modifications do not necessarily alter the current most when centered in the k-mer, and using the prior analysis allowed picking the correct position for each modification to conduct the quantification (Figure 2C). Tombo provides quantification of RNA modifications by comparing current levels between the *E. coli* rRNA with those found in the synthetic RNA without modifications.^26^ The base call data derived from the current levels can also be analyzed for RNA modification quantification. The computational tools ELIGOS2 and Nanopore-Psu were employed. The tool ELIGOS2 can be used for modification quantification by comparing the error at specific bases (ESB) for the *E. coli* rRNA strands against the synthetic rRNA by a method previously described;^13,34^ Nanopore-Psu was trained on mostly cellular rRNA to predict Ψ occupancy specifically in nanopore sequenced RNA.^14^

**Figure 3.**
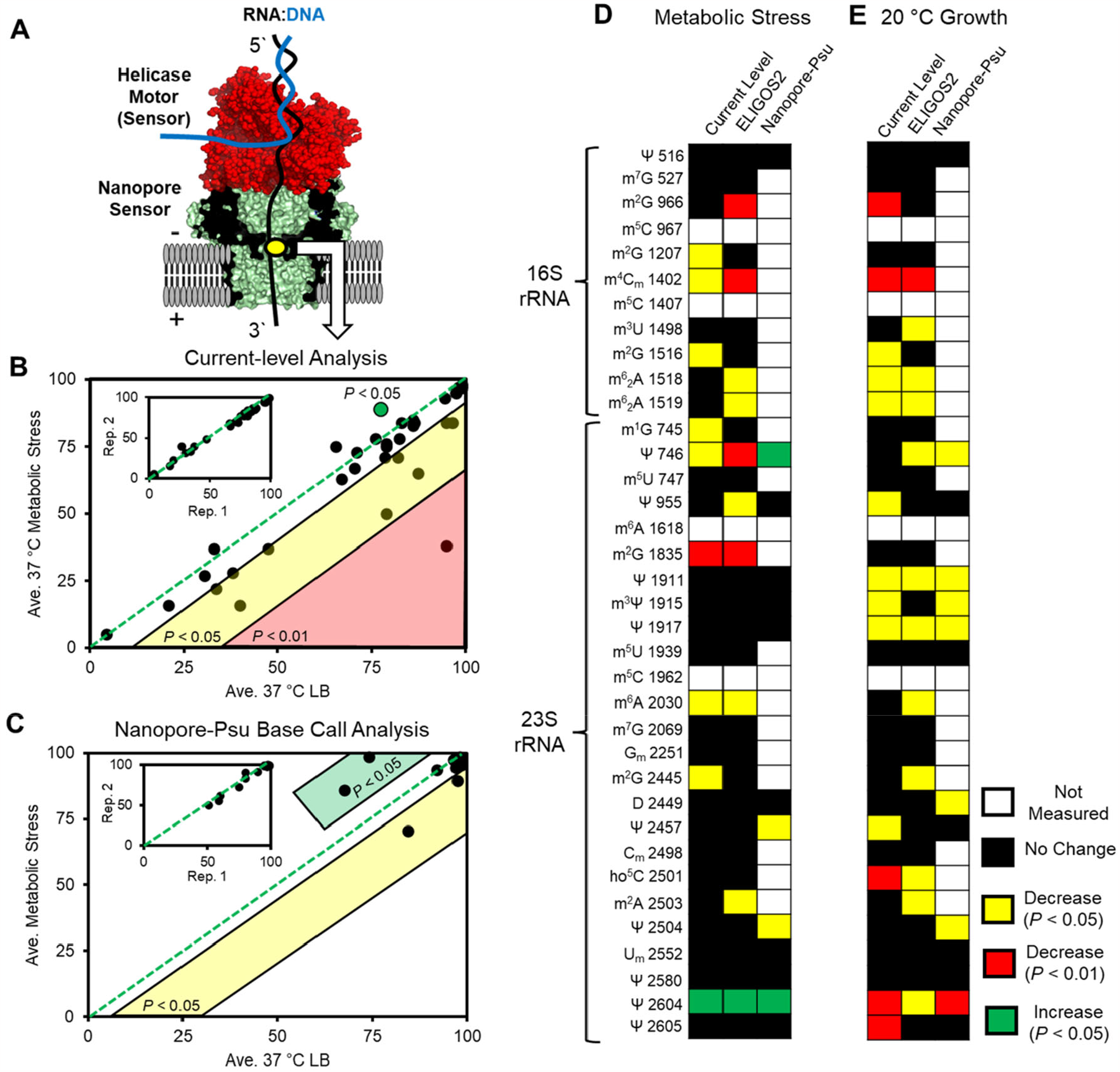
Stress-dependent changes in *E. coli* rRNA modifications found by direct nanopore RNA sequencing from the nanopore protein sensor. (A) The nanopore sequencing setup has two sensors, the helicase and protein nanopore, in which the nanopore provides current levels used in base calling analysis. (B) Current-level quantification via the Tombo tool for changes in rRNA modifications. Inset illustrates biological replicate reproducibility. (C) Base calling quantification for Ψ sites using Nanopore-Psu. Inset shows reproducibility in biological replicates. Changes in the *E. coli* rRNA modifications identified after (D) metabolic or (E) cold-shock stress. The changes were color-coded based on a Student’s t-test of the RNA modification level changes between normal growth and stress conditions (yellow = decrease with *P* < 0.05; red = decrease with *P* < 0.01; and green = increase with *P* < 0.05).

Biological replicates for sequencing *E. coli* rRNA obtained from cells grown in LB media at 37 °C found the modification quantification from current-level and base-call data were generally reproducible (Figures 3B and 3C inset). Next, scatter plot analysis of the RNA modification levels under normal conditions vs. the stress conditions identified many sites that deviated between them (Figures 3B, 3C, and SX). Duplicate data sets were analyzed for each condition and the Student’s t-test was used to find those with significant changes; the changes were color-coded to be yellow for a lower level of significance in the decrease (\**P* < 0.05), red for a larger significance in the decrease (\*\**P* < 0.01), and green for a lower level of significance in an increase in the change (\**P* < 0.05; Figures 3D and 3E).

Metabolic stress to the *E. coli* was induced by growing them on minimal media (M9 with 0.5% glucose) under aerobic conditions at 37 °C for 20 h following literature reports.^21^ Cold-shock stress to the *E. coli* was induced by first growing the cells aerobically in LB media at 37 °C for 20 h, followed by placing the cells at 20 °C and then allowing them to grow for 1 h under the changed temperature. Heat-shock stress was studied and the data will be provided a peer-reviewed publication with the knowledge that the RNA sequenced was fragmented from the high temperature, a known phenomenon.^35^ The cells exposed to metabolic stress showed significant decreases in the rRNA modifications based on current-level analysis (8 sites), ELIGOS2 base-call analysis (9 sites), and Nanopore-Psu base-call analysis (2 sites; Figure 3D). In all three data analysis approaches, the 23S Ψ2604 was found to increase in occupancy (Figure 3D). The analysis provided contradictory findings for 23S Ψ746, in which a decrease in occupancy was found with Tombo and ELIGOS2 but an increase in occupancy was found for Nanopore-Psu (Figure 3D). Cold-shock stress to *E. coli* resulted in rRNA modification level decreases based on current-level analysis (13 sites), ELIGOS2 analysis (12 sites), and Nanopore-Psu analysis (7 sites; Figure 3E).

The rRNA modification levels found to change with metabolic or cold-shock stress can be grouped. The first group is m^2^G, in which the rRNAs have 5 sites of this modification (Figure 1B), four of which decreased with metabolic stress, and three decreased with cold-shock stress (Figures 3D and 3E). The second group is Ψ, in which *E. coli* rRNAs have 11 sites (10 Ψ sites and 1 m^3^Ψ site; Figure 1B), three changed with metabolic stress, and nine changed with cold-shock stress (Figures 3D and 3E). The third group is a few sites of single modification structures that changed and include cold shock resulting in a decrease of 16S m^4^C_m_1402, 16S m^6^_2_A at positions 1518 and 1519, 23S m^6^A2030, 23S ho^5^C2501, and 23S m^2^A2503 (Figure 3E). Metabolic stress resulted in a decrease at 16S m^4^C_m_1402, the two 16S m^6^_2_A sites at positions 1518 and 1519, and 23S m^2^A2503 (Figure 3D). These findings identify rRNA modifications can change with metabolic and thermal stress in *E. coli*. This is a finding in line with a hypothesis that rRNA play a central role in sensing stress.^36^

### The helicase sensor identifies the methyl group lost from m^4^C_m_ during stress

The hypermodified RNA m^4^C_m_ stands out as one for further investigation because its levels decrease with both forms of stress studied. Does stress cause loss of both the sugar and base methyl groups, or only one methyl group? And if it is just one, which one is it (Figure 4A)? The 16S m^4^C_m_1402 is the only modification of this type in *E. coli*, and studying its change by mass spectrometry is very challenging; further, if only one methyl group is lost, analyzing rRNA fragments by mass spectrometry cannot easily differentiate whether it was the base or sugar methyl group. The helicase sensor was found to provide a solution to this challenge using the same analysis we previously reported for finding Ψ in the SARS-CoV-2 RNA genome (Figure 4B).^12^

**Figure 4.**
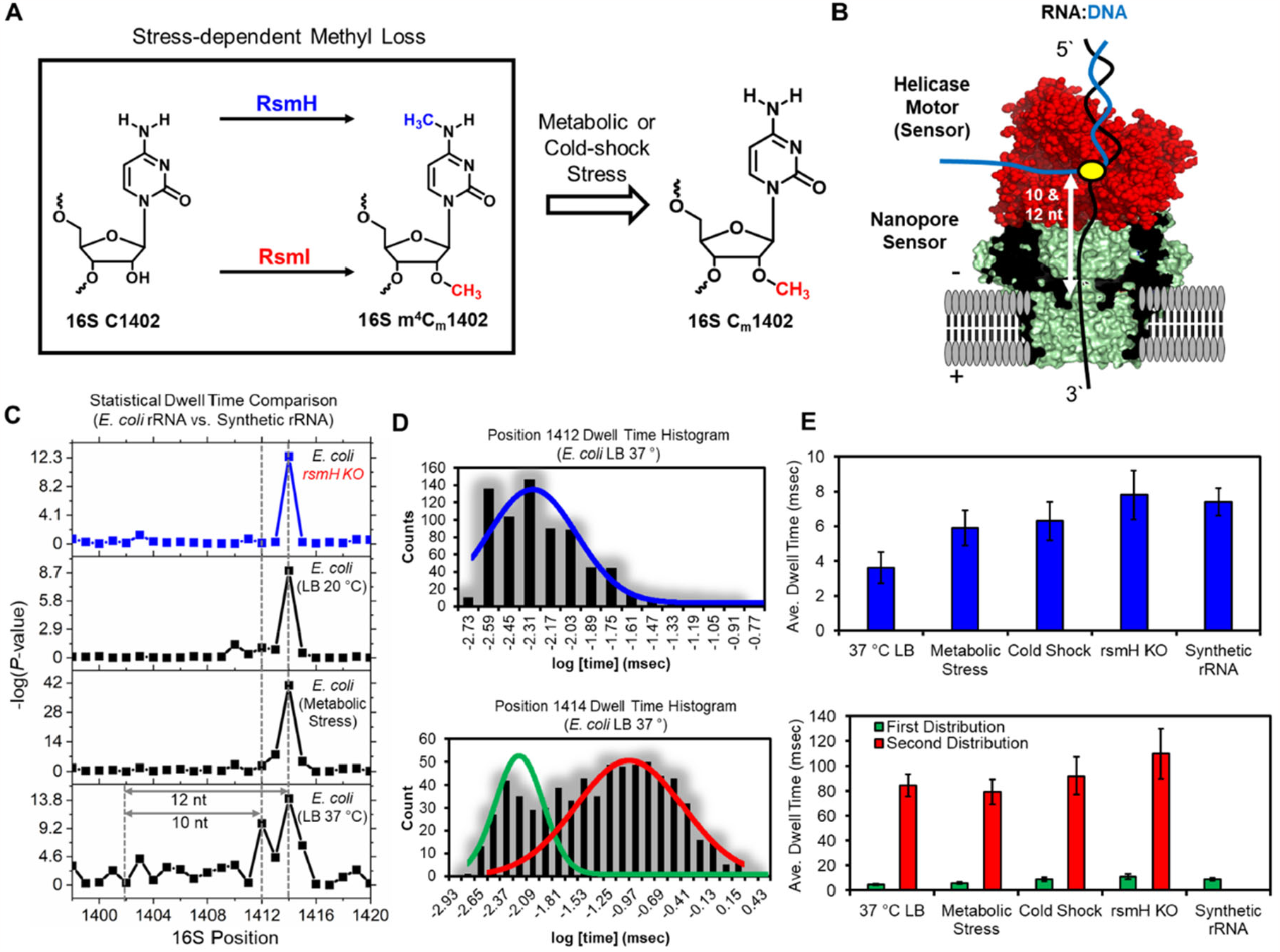
The helicase sensor allows identification of the methyl group lost from the *E. coli* 16S m^4^C_m_1402 site during metabolic or cold-shock stress. (A) The analysis provided in Figures 3D and 3E found that the bis-methylated base 16S m^4^C_m_1402 had changed. Were one or two methyl groups lost, and if one, which one? (B) The helicase sensor identifies RNA modifications by a change in dwell time; however, as a result of the base calling coming from ionic current levels in the nanopore sensor, the modifications are observed in the reference transcriptome alignment as a change in dwell time that is 3′ distal by 10-12 nucleotides from the modification position in the RNA. (C) Nanocompore analysis was used to compare two biological replicates of the *E. coli* 16S rRNA vs. the synthetic rRNA without modifications to find dwell times that differ significantly between the samples. The P-values reported from Nanocompore were negative-log transformed for visualization, in which an increase in value represents a greater degree of difference between the samples. (D) Histograms of dwell time values obtained from Nanopolish analysis of 16S rRNA at positions 1412 and 1414 from non-stressed *E. coli*. (E) Average dwell times were found for the rRNAs being unwound by the helicase sensor by fitting the dwell time histograms to a Gaussian distribution (panel D). The errors reported are for the fitting errors of the average value plotted.

The Nanocompore tool was used to identify statistically significant differences (Kolmogorov-Smirnov (KS) test) in the helicase sensor dwell times between the *E. coli* 16S rRNAs and the synthetic 16S rRNA without any modifications.^15^ The *P*-values from the KS test were negative-log transformed for visualization purposes such that an increase in value represents a greater degree of statistical significance between the datasets. The statistical analysis of *E. coli* grown at 37 °C in LB media identified two positions (1412 and 1414) for which 16S m^4^C_m_1402 impacted the helicase dwell time (Figure 4C bottom panel). We hypothesize one is for the base methyl group and the other is the sugar methylation. Surprisingly, the metabolic and cold-shock stress experiencing *E. coli* rRNA when statistically analyzed against the control were found to have an altered dwell time at position 1414 supporting one methyl group was lost from the same site under both stressors (Figure 4C interior panels). Confirmation that it was the loss of the base methyl group was derived from sequencing the 16S rRNA from *E. coli* in which the methyltransferase for writing the methyl group on *N*^*4*^ of C 1402 was knocked out (rsmH). Analysis of the rsmH knockout *E. coli* 16S rRNA found only the 1414 site had a significant difference in dwell time compared to the synthetic rRNA (Figure 4C top panel). The analysis identifies in *E. coli* that metabolic and cold-shock stress results in the 16S rRNA having a C_m_ residue at position 1402 instead of a hypermodified m^4^C_m_. This demonstration provides an example of the power of the helicase as a sensor to learn details about RNA modifications. This would not be easily achieved by other methods.

To gain a better biophysical understanding of how the m^4^C_m_ modification was changing the helicase activity (i.e., dwell time), hundreds of events were analyzed with Nanopolish to measure the dwell time values at positions 1412 and 1414 in the samples.^25^ The dwell time values were log-transformed and then binned and histograms were made followed by fitting with a Gaussian distribution function (Figure 4D). At position 1412 for the *N*^*4*^ methyl group on C, the distribution was fit to a single Gaussian function, and at position 1414 for the 2′-*O*-methyl group on C, the distribution was fit to two Gaussian functions. The base methyl group impacting position 16S 1412 was found in the modified 37 °C grown *E. coli* to have a dwell time average of ∼3 msec; loss of the methyl group in the stressed *E. coli, rsmH* KO cells, and synthetic 16S rRNA gave an average dwell time of nearly 2-fold or longer (6-8 msec). This analysis identifies the presence of the base methyl group results in faster helicase activity compared to the canonical nucleotide.

Inspection of the data at position 1414 found two populations in all the *E. coli* rRNAs (no stress, stress, and *rsmH* gene knockout) with an average dwell time of less than 10 msec and a population with an average dwell time increased by about 10-fold (80-110 msec; Figure 4E). The most revealing observation is that in the synthetic 16S rRNA at position 1414 only one population of helicase dwell times was observed with an average dwell time of ∼9 msec (Figure 4E). This final analysis identifies the 2′-*O* methyl group on C, impacts helicase processivity with two different time values, one that is 10-fold longer than observed for the unmodified C nucleotide and the other with a similar time value as measured for the C nucleotide.

### Identification of *E. coli* rRNA operons and their deposition of the 11 Ψ residues

In the final analysis, the rRNA sequencing reads were aligned against the *E. coli* reference genome with all seven rRNA operons (*rrnA, B, C, D, E, G*, and *H*; Figure 5A) instead of the *rrnA* operon reference in the previous studies. The operons each possess their own sequence variations that allow alignment to each operon (Figure 5B).^21^ One drawback to this is that the data at each site have lower coverage, and therefore, the operon-specific analysis used Nanopore-Psu focused on 10 Ψ sites and 1 m^3^Ψ in the *E. coli* rRNA sequences. As a consequence of Ψ being the most abundant modification in the rRNA and these modifications being subject to change, the limited analysis provides a picture of operon-specific modification changes before and after stress. In Figure 5, the analysis is highlighted for metabolic stress. The operon-specific analysis of cold-shock stress will be provided in a peer-reviewed publication with the limitation that not all sites could be analyzed as a result of the decreased sequencing depth when aligning to all seven operons.

**Figure 5.**
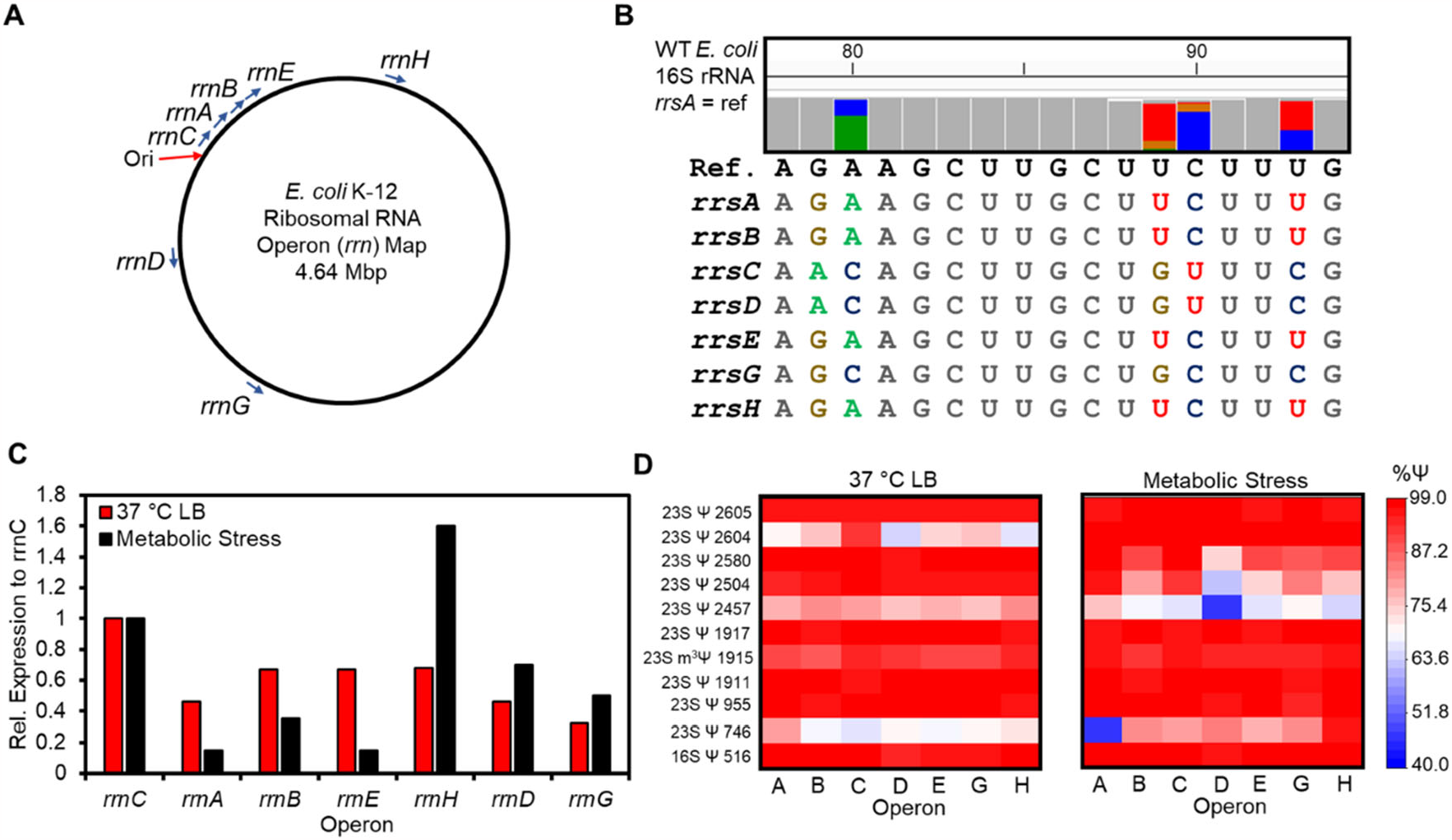
Operon-specific rRNA changes in Ψ writing in *E. coli* rRNAs. (A) Genome map illustrating the positions of the rRNA operons in the E. coli genome. (B) Example IGV plot of *E. coli* 16S rRNA from positions 78-93 showing five different natural operon-specific sequence variations. (C) Relative expression levels for the seven different *E. coli* rRNA operons when the cells were grown aerobically at 37 °C in either LB media or under metabolic stress. (D) Heat plots for Nanopore-Psu predicted levels for the 11 Ψ sites in the 16S and 23S rRNAs. The values plotted are averages from a set of biological replicates.

First, the expression levels of rRNA from each operon were determined under normal and metabolic stress conditions. A prior report normalized the data to the expression of *rrnC* because it is the closest to the origin of replication (Ori; Figure 5A),^21^ and we maintain that convention herein. When *E. coli* were grown in LB media at 37 °C, the most highly expressed operon was *rrnC* followed by *rrnB, rrnE, rrnH, rrnA, rrnD*, and *rrnG* (Figure 5C red bars). In general, the highly expressed operons were those in the direction of replication (*rrnC, B, E, H*, and *A*), and the two operons oriented in the opposite direction were expressed at lower levels (Figures 5A and 5C). Next, we found that under metabolic stress, the operon expression levels changed (Figure 5C). Those more distal to the Ori (*rrnH, D*, and *G*) showed the greatest increase in expression while those more proximal to the Ori (*rrnA, rrnB*, and *rrnE*) decreased in expression (Figures 5A and 5C). A previous report conducted a similar analysis to that described here,^21^ by sequencing the *E. coli* rRNAs via conversation to cDNAs and they obtained nearly identical results as found from direct nanopore RNA sequencing conducted herein.

The Nanopore-Psu analysis for Ψ in each of the rRNAs from the seven operons is the first experiment of this type and provides deeper insight into the changes found in Figure 3. Pseudouridine levels in the rRNAs under normal growth conditions, in general, were similar across all seven rRNA operons, with the exceptions of 23S Ψ746 and 23S Ψ2604 (Figure 5D left panel). The installation of 23S Ψ746 was predicted by Nanopore-Psu to exist with the highest frequency of 80% in the rRNA from the *rrnA* operon, and at the lowest frequency of 67% in the rRNA from the *rrnC* operon. In contrast, 23S Ψ2604 was installed with the highest frequency in the rRNA from the *rrnC* operon at 93% and the lowest frequency in the rRNA from the *rrnD* operon at 65%.

Under metabolic stress, operon-specific Ψ levels were similar at eight sites with three noteworthy differences. The first is 23S Ψ 2604, in which the installation increased the most on all operons except *rrnC*, which was already at a high level as described above. When the Ψ levels were determined by alignment to a single reference (Figure 3E), 23S Ψ2604 was found to increase under metabolic stress; however, the operon-specific analysis finds this increase occurs on all operons except *rrnC* (Figure 5D). The second is 23S Ψ2457, which decreased in installation in the rRNA when aligning the data against a single reference (Figure 3E), and this claim holds when aligning the reads against all seven operons (Figure 5E right panel). An interesting point is that the decrease in writing 23S Ψ2457 occurred in the *rrnD* operon. The second is 23S Ψ746, which was found in the single operon alignment data to not change upon metabolic stress; in contrast, alignment of the data to all seven operons finds this Ψ increases in most operons and decreases in *rrnH* leading to the null result in the single operon alignment data (Figures 5E right panel and 3E). This final observation identifies stress can result in underlying rRNA modification changes with operon specificity that are not observed when aligning the data to a single reference.

## Discussion

### Direct RNA nanopore sequencing can monitor 16 different *E. coli* rRNA modifications

In the present work, *E. coli* rRNA was sequenced directly with the commercial nanopore platform from ONT. This was done to evaluate the ability to sequence and quantify up to 17 different RNA modification structures that occur 36 times in these RNAs. Prior structural and in cellulo studies have determined the sites, quantified the installation, and identified the writers for all but one of these 36 modifications in *E. coli* rRNA;^2,20,28^ this provides an excellent case to study direct RNA nanopore sequencing for more than one modification type at a time during sequencing. Finally, a prior study reported on two modifications in the 16S *E. coli* rRNA (m^7^G527 and Ψ516), and the present work that expands the analysis to all rRNAs is fully consistent with the earlier report on these two sites.^11^

Seventeen of the 36 modifications are located in the primary sequence distant from other rRNA modifications, and they were easily evaluated to determine their ionic current, dwell time, and/or base calling signatures. All of these could be revealed in one or more of the nanopore sequencing data types with the exception of m^5^C. This observation is consistent with a prior report on mRNA sequencing.^13^ Another observation is with respect to the two m^6^A sites in the 23S rRNA (positions 1618 and 2030), in which 1618 was only observable by a helicase dwell signature while 2030 yielded a base call and dwell signature. Computational tools such as Epinano^37^ and ELIGOS2^13^ that inspect base call data can locate and quantify m^6^A in mRNA, in which this modification is deposited at high frequency in DR**A**CH motifs (D = A or G; R = A or G; H = A, C, or U; bold = m^6^A). In contrast in *E. coli* rRNA, 1618 and 2030 are written in the sequence contexts 5′-AC**A**GG and 5′-UG**A**AG, respectively. This observation nicely demonstrates the strong sequence context dependency in direct RNA nanopore sequencing signatures, a feature we previously described for Ψ.^34^

Nineteen of the 36 RNA modifications to the *E. coli* rRNA are present as clusters in the primary sequence representing a challenge to deconvolute the nanopore and helicase signatures.

We approached this challenge by sequencing rRNA strands from *E. coli* that had known writer enzymes knocked out to identify the signal changes in comparison to the rRNAs from the native *E. coli* cells. This approach worked with two exceptions that include the two adjacent m^6^_2_A sites at 1518 and 1519 in the 16S rRNA, and the three closely spaced 23S Ψ1911, m^3^Ψ1915, and Ψ1917 sites; monitoring these was achievable in the base call data (Figure 2E). A key success in this approach for determining the modification signatures was the four closely spaced modifications in the 23S rRNA (C_m_2498, ho^5^C2501, m^2^A2503, and Ψ2504). As a final point, synthetic RNAs made by IVT provided independent verification of the signatures for m^5^C, m^2^G, C_m_, m^4^C, ho^5^C, and m^5^U.

### Acknowledgment of limitations to the data analysis

The significant prior studies on *E. coli* rRNA allow comparison of the nanopore data reported in the present work.^2,20,28^ With respect to RNA modification quantification, a convenient aspect of nanopore data is the ability to predict modification occupancy. Quantification of the data is successful, as reported here and in other studies, but these values when compared to high-accuracy mass spectrometry data are quite variable. For example, m^7^G (16S 527 and 23S 2069) is quantifiable by the current level to give values similar to MS measured values (16S m^7^G527: nanopore = 98% and MS >95%;^20^ Figure SX). In a less ideal case, 23S G_m_ 2251 is present at >95% occupancy by MS analysis,^20^ while the current-level analysis predicts 28% occupancy, base call analysis fails to make a prediction (Figure SX), and helicase analysis is challenging to quantify, as described below. Similar large differences between MS and nanopore values exists for m^6^A, m^5^C, m^5^U, D, and ho^5^C, as well as a single m^2^G site (23S 2445). Direct RNA nanopore sequencing can provide data not obtainable by MS analysis, as demonstrated by the two Ψ sites at positions 2604 and 2605 in the 23S rRNA. In this final example, MS analysis is challenged to quantify the modifications,^30^ while base call data and ionic current level data can be quantified, albeit they give different results (base call: 2604 = 72% and 2605 = 98%; current level: 2604 = 50% and 2605 = 55%). These comparisons indicate nanopore data can be quantified but the values may differ from more accurate MS data. For this reason, we looked for changes in values instead of absolute quantification of occupancy.

Another noteworthy limitation is in regards to using base calling as a means of RNA modification detection and quantification. The seven *E. coli* rRNA operons serve as an excellent example of caution in this analysis. For instance, in the 16S rRNA at position 93 there exists a natural C/U sequence variation between the operons (Figure 5B). Interestingly, a C miscall is the most common Ψ signature in nanopore base call data;^10,12,16,34^ thus, this natural sequence variation masquerades as a Ψ if only base call data is considered, when in actual fact one is sequencing a mixture of C and U variants at that site. This example nicely illustrates why in our prior studies the use of helicase dwell time, for which Ψ yields a signature, serves as a check for verification when using direct RNA nanopore sequencing for the discovery of new Ψ sites.^12^

### Stress-dependent rRNA changes in *E. coli* monitored by nanopore sequencing

Using the analysis of rRNA modifications just described, the *E. coli* cells were subjected to either micronutrient or cold-shock stress followed by reanalysis of the rRNA modifications. Keeping the quantification challenges in mind, the data were analyzed for changes in the rRNA modifications relative to the cells grown under normal conditions (Figure 3). When the *E. coli* cells were allowed to grow and divide under either stress condition, the modifications did not change or were found to decrease in occupancy with the except of one that increased in occupancy. Modifications that consistently decreased with the stress based on a change in one of the data types were m^2^G (16S 1516 and 23S 2445) and Ψ (23S 746, 955, 2457, and 2504).

There are two exceptions to this observation; under metabolic stress, 23S Ψ2604 was found to increase in its occupancy, and mixed results were found for 23S Ψ746 (current and ELIGOS2 predict a decrease, and Nanopore-Psu predicts an increase; Figure 3D). Clarification of the inconsistent result for 23S Ψ746 comes from the operon-specific Ψ analysis (Figure 5D). This position was found to have mixed operon-specific Ψ deposition, in which the *rrnA* operon had less Ψ installed with metabolic stress, and the others showed an increase in Ψ installation.

The data in Figure 3 were obtained by aligning to only the *rrnA* operon as a reference; hence, the alignment is likely the culprit. For the current-level analysis with Tombo and base call analysis with ELIGOS2, minimap2 was used with one set of parameters for alignment, while Nanopore-Psu conducts the alignment with minimap2 using a different set of parameters; this difference likely impacted the reads mapped, and the Ψ quantified. This demonstrates how small differences in computational tools can impact the results of the analysis.

### Helicase sensor dwell-time analysis provides clarity on the m^4^C_m_ change with stress

In the 16S rRNA sequence at position 1402, there is a hypermodified C nucleotide m^4^C_m_ that decreased with metabolic and cold-shock stress (Figures 2D and 2E). First, there exists a m^5^C at position 1407 in this sequence (Figure 1B), but as described, this modification does not impact nanopore data, and therefore, the signature results from m^4^C_m._ What was the change, loss of one or both methyl groups? If one, which methyl group was lost? In *E. coli*, this is the only m^4^C_m_, and to study this by complete digestion to nucleosides followed by quantitative MS analysis would be challenging, especially because *E. coli* have many C_m_ and m^4^C residues in other RNAs; moreover, using MS to analyze rRNA fragments would fail to provide a conclusive result with loss of a single methyl group.

The helicase sensor was found to provide data to answer this question. The hypermodification m^4^C_m_ impacts the helicase activity in two different positions that are found when comparing helicase dwell time differences between the wild-type *E. coli* rRNA with the synthetic rRNA without modifications. A prior study noted that this residue could impact the helicase dwell time,^18^ consistent with our findings. Under the stressors imposed, only one helicase dwell time difference was observed in the statistical comparison (Figure 4C). The deduction of which methyl group was lost came from studying rRNA from *E. coli* with the base methyl writer genes *rsmH* knocked out. The comparative analysis found that when only a C_m_ resides at position 1402, a single helicase dwell time difference is observed that tracks with those observed in the stressed *E. coli* (Figure 4C). This confirms under stress in the 16S rRNA at position 1402, the cells only methylate the sugar to yield C_m_.

In principle, a histogram for a population of dwell times that differ between the modified and unmodified states can be integrated to quantify the presence of modification. The histogram shown in Figure 4D illustrates a challenge with the quantification of helicase dwell time. The nanopore is a single-molecule sensor, and the dwell times represent monitoring helicase activity one molecule at a time. At this molecular level, the data identify how stochastic enzymes are in processing their substrates, resulting in the broad time distributions measured.^38^ Therefore, any unmodified RNA may be masked by the presence of the modification in the helicase dwell time resulting in less reliable quantification. Nonetheless, the presence of the dwell time change provides a check for the presence of an RNA modification when one exists. This feature may also exist when sequencing DNA modifications with the nanopore.^39^

Two remaining questions regarding the helicase activity on m^4^C_m_ include these: why are there two different points at which this double modification changes the helicase activity? Secondly, why does C_m_ yield two different helicase dwell times? The patent literature suggests the helicase used by ONT is a mutated version of a Hel308 helicase,^40^ but the actual mutations are not publicly known. With this limitation and using the native structure for archaeal Hel308 solved bound to DNA, we can hypothesize why a base and sugar methyl group would impact the helicase at different positions.^41^ In this structure, Lys 289 interacts with the ribose backbone impacting 2′-*O*-methyl group, while the base interacts with Phe 350. Interestingly, these two interactions occur two nucleotides apart, consistent with the nanopore helicase dwell time analysis for m^4^C_m_. A reason for two helicase dwell time populations (Figure 4D) may result from C_m_ favorably adopting 3’-*endo* over the 2’-*endo* sugar pucker, in which the 3’-*endo* conformation likely stabilizes RNA duplexes by preforming the A helix and the 2’-*endo* does not;^42^ however, studies to understand the kinetics of this equilibrium in a helicase active site have yet to be conducted to support whether they are sufficiently long-lived to impact the helicase activity differently.

### Biological implications for rRNA modification changes during stress

The implications of rRNA modification changes in *E. coli* under metabolic or cold-shock stress are described for a few of those found in this study. The first is the decrease of 16S m^4^C_m_ 1402 yielding C_m_ at this position. This residue is conserved in bacteria and is located in the P-site of the ribosome where it makes contact via the *N*^*4*^ nitrogen with the second and third positions of the P-site codon.^43^ The function of this contact is proposed to fine-tune the P-site structure resulting in correct recognition of the initiation codon. Loss of the *N*^*4*^ methyl group from 16S m^4^C_m_1402 results in increased efficiency of non-AUG start codons and greater UGA read-through, resulting in decreased ribosome fidelity.^43^ This may be a feature used by the cell to assist survival during stress.

A second noteworthy rRNA modification that decreased under cold-shock stress is the 23S ho^5^C2501 (Figure 2E). This site is of interest to us because it is the only modification resulting from a two-electron oxidation of the base. This modification site is located in proximity to the peptidyl transferase center and is hydroxylated in most bacteria.^44^ The presence of this modification is hypothesized to provide tolerance to oxidative stress;^44^ however, why it is lost during cold-shock stress remains unknown. Two possibilities include, (1) transferring *E. coli* at stationary phase growth at 37 °C to 20 °C leads to new rRNA synthesis and because this is a late-stage modification in the 23S rRNA,^44^ it was not yet installed, resulting in the lower yields. (2) ho^5^C is easily further oxidized, and cold-shock stress could have resulted in conditions that led to its further oxidation yielding a highly-distorted base compared to the parent.^45^ Signals were not observed that would suggest the presence of the distorted structure, but it is possible this structure could not easily pass the helicase, and therefore, was not detected in the longer reads. Future work is needed to understand why 23S ho^5^C2501 changed.

The changes in Ψ observed with stress are not easily understood from a structural perspective. Loss of any Ψ in *E. coli* rRNA has no or a small phenotype; actually, they can all be removed and the cells only show modest defects in ribosome biogenesis, function, and cell growth.^46^ As for metabolic stress, prior proteome analysis provides details about changes in protein writers during stress that can be tied to the changes observed in the sequencing experiments in the present work.^47^ An increase in 23S Ψ2604 occurs with metabolic stress, and the expression level of the writer for this Ψ (rluF) increases by 150%. Furthermore, rluF installs Ψ in tRNA^Tyr^ at position 35, and this could also be impacted by metabolic stress. Stress-dependent changes in tRNA modifications have been noted.^48^

In contrast, lower levels of 23S m^6^A2030, m^2^G2445, and Ψ2504 were observed that track with the reduction in expression of the corresponding writers (rlmJ, rlmL, and rluC, respectively)^47^ for these rRNA modifications during stress. The most significant was 23S m^2^G1835, which gave a large decrease in occupancy in the rRNA, and the proteome analysis failed to identify the presence of the writer protein rlmG in *E. coli* experiencing the same form of metabolic stress as imposed in these studies.^47^ This argument for the RNA modification changes is not the complete picture, and future studies likely assisted with direct RNA nanopore sequencing will be needed to further address the biology of the changes. The changes in *E. coli* rRNA modifications under stress differ from those reported for *S. cerevisiae* found by direct RNA nanopore sequencing.^10^ The eukaryotic vs. prokaryotic cell types studied and the difference in the time of the imposed stress relative to the cell replication rate best address the differing findings. These comparisons to prior studies support a conclusion in *E. coli* that rRNA and its modification status are sensitive to environmental conditions. These rRNA modifications may play a larger role in cell signaling beyond supporting ribosome structure for protein translation. The present work now provides nanopore current levels, base call information, and helicase dwell time analyses for 10 additional RNA modifications (m^4^C_m_, m^2^G, m^5^U, m^3^U, m^1^G, m^3^Ψ, D, ho^5^C, m^2^A, and m^4^C). These details will assist future direct RNA nanopore sequencing experiments for discovery of how many more RNA modifications, beyond those already known, are impacted when cells are exposed to environmental challenges.

## Methods

### *E. coli* growth conditions

The *E. coli* K12 DH5 alpha strain with an amp-resistant gene supplied from a plasmid (psiCheck2) was grown aerobically in ampicillin-containing LB media at 37 °C in a shaking incubator to stationary phase conditions (∼20 h). The knockout *E. coli* K12 strains obtained from the Keio collection are kanamycin resistant and therefore, were grown aerobically in kanamycin-containing LB media in a shaking incubator at 37 °C to stationary phase. Induction of metabolic stress was achieved following the literature,^21^ in which the wild-type *E. coli* were grown in minimal media (M9) with 0.5% glucose to the stationary phase. The thermal stress was induced following the literature by taking the *E. coli* grown for 15 h and then they were mixed 1:1 with media at either 20 °C or 42 °C and then grown for 1 h under the new temperature. A colony-forming unit assay was conducted on all *E. coli* samples to verify the cells were still viable.

### RNA extraction

After the growth, the cells were pelleted by centrifugation at 6800 rpm for 10 min and the supernatant was decanted. The cell pellet was then mixed with 1.8 mL of TRIzol reagent (Life Technologies) and lightly vortexed to resuspend the cells in the reagent. Following the instructions for the Zymo Direct-zol RNA Miniprep kit, the total RNA was purified away from the other cellular components. The RNA was verified by agarose gel electrophoresis and then stored at -80 °C until sequencing was commenced.

### Preparation of synthetic RNA

The duplex DNA utilized for the in vitro transcription of the 16S and 23S sequences found in the *rrnA E. coli* operon, and the DNA for the judiciously designed RNAs to study individual modifications were obtained from a commercial source (sequences are provided below; Twist Biosciences and Azenta Life Sciences). The DNA for IVT used a T7 RNA polymerase promoter. The RNAs were synthesized using the T7 MegaScript Kit (Thermo Fisher Scientific) following the manufacturer’s protocol. The modified RNA strands were prepared by IVT by substituting the canonical NTP with the modified NTP. The following NTPs were obtained from Trilink Biotechnologies with the reported purities in parentheses m^4^C (>90%), C_m_ (>98%), ho^5^C (>95%), or m^5^C (>95%). The NTPs used for the synthesis of RNAs containing m^5^U, m^2^G, and m^4^C_m_ were synthesized and characterized as described below. The modified nucleotides were installed in the RNA by replacing the canonical form with the modified form at the same concentration stated by the IVT kit. The one exception was the RNA synthesized with m^2^G. Because the first nucleotide inserted by T7 RNA polymerase to achieve high-efficiency synthesis is a G that was found to be stalled by m^2^G, the reaction was doped with 10 mM GMP to initiate synthesis while all other sites had the m^2^GTP installed. Success in the synthesis of the RNA was determined by agarose gel electrophoresis. Because we were able to synthesize RNA strands with either m^4^C or C_m_ by IVT, an attempt to make an RNA with m^4^C_m_ was conducted; however, after repeated trials, no full-length RNA was made and therefore the synthesis was not further pursued.

### Sequences for the coding strand of duplex DNA used for the synthesis of RNA

The bold sequences are the T7 promoter on the 5’ end and the poly-A tail for library preparation on the 3’ end.

#### 16S (rrsA)

##### 5’-AAGCTAATACGACTCACTATAGG

AAATTGAAGAGTTTGATCATGGCTCAGATTGAACGCTGGCGGCAGGCCTAACACATGCAAGTCGAACGGT

AACAGGAAGAAGCTTGCTTCTTTGCTGACGAGTGGCGGACGGGTGAGTAATGTCTGGGAAACTGCCTGAT

GGAGGGGGATAACTACTGGAAACGGTAGCTAATACCGCATAACGTCGCAAGACCAAAGAGGGGTACCTTC

GGGCCTCTTGCCATCGGATGTGCCCAGATGGGATTAGCTAGTAGGTGGGGTAACGGCTCACCTAGGCGAC

GATCCCTAGCTGGTCTGAGAGGATGACCAGCCACACTGGAACTGAGACACGGTCCAGACTCCTACGGGAG

GCAGCAGTGGGGAATATTGCACAATGGGCGCAAGCCTGATGCAGCCATGCCGCGTGTATGAAGAAGGCCT

TCGGGTTGTAAAGTACTTTCAGCGGGGAGGAAGGGAGTAAAGTTAATACCTTTGCTCATTGACGTTACCC

GCAGAAGAAGCACCGGCTAACTCCGTGCCAGCAGCCGCGGTAATACGGAGGGTGCAAGCGTTAATCGGAA

TTACTGGGCGTAAAGCGCACGCAGGCGGTTTGTTAAGTCAGATGTGAAATCCCCGGGCTCAACCTGGGAA

CTGCATCTGATACTGGCAAGCTTGAGTCTCGTAGAGGGGGGTAGAATTCCAGGTGTAGCGGTGAAATGCG

TAGAGATCTGGAGGAATACCGGTGGCGAAGGCGGCCCCCTGGACGAAGACTGACGCTCAGGTGCGAAAGC

GTGGGGAGCAAACAGGATTAGATACCCTGGTAGTCCACGCCGTAAACGATGTCGACTTGGAGGTTGTGCC

CTTGAGGCGTGGCTTCCGGAGCTAACGCGTTAAGTCGACCGCCTGGGGAGTACGGCCGCAAGGTTAAAAC

TCAAATGAATTGACGGGGGCCCGCACAAGCGGTGGAGCATGTGGTTTAATTCGATGCAACGCGAAGAACC

TTACCTGGTCTTGACATCCACGGAAGTTTTCAGAGATGAGAATGTGCCTTCGGGAACCGTGAGACAGGTG

CTGCATGGCTGTCGTCAGCTCGTGTTGTGAAATGTTGGGTTAAGTCCCGCAACGAGCGCAACCCTTATCC

TTTGTTGCCAGCGGTCCGGCCGGGAACTCAAAGGAGACTGCCAGTGATAAACTGGAGGAAGGTGGGGATG

ACGTCAAGTCATCATGGCCCTTACGACCAGGGCTACACACGTGCTACAATGGCGCATACAAAGAGAAGCG

ACCTCGCGAGAGCAAGCGGACCTCATAAAGTGCGTCGTAGTCCGGATTGGAGTCTGCAACTCGACTCCAT

GAAGTCGGAATCGCTAGTAATCGTGGATCAGAATGCCACGGTGAATACGTTCCCGGGCCTTGTACACACC

GCCCGTCACACCATGGGAGTGGGTTGCAAAAGAAGTAGGTAGCTTAACCTTCGGGAGGGCGCTTACCACT

TTGTGATTCATGACTGGGGTGAAGTCGTAACAAGGTAACCGTAGGGGAACCTGCGGTTGGATCACCTCCT

TA **AAAAAAAAAAAA**

#### 23S (rrlA)

##### 5’-AAGCTAATACGACTCACTATAGG

GGTTAAGCGACTAAGCGTACACGGTGGATGCCCTGGCAGTCAGAGGCGATGAAGGACGTGCTAATCTGCG

ATAAGCGTCGGTAAGGTGATATGAACCGTTATAACCGGCGATTTCCGAATGGGGAAACCCAGTGTGTTTC

GACACACTATCATTAACTGAATCCATAGGTTAATGAGGCGAACCGGGGGAACTGAAACATCTAAGTACCC

CGAGGAAAAGAAATCAACCGAGATTCCCCCAGTAGCGGCGAGCGAACGGGGAGCAGCCCAGAGCCTGAAT

CAGTGTGTGTGTTAGTGGAAGCGTCTGGAAAGGCGTGCGATACAGGGTGACAGCCCCGTACACAAAAATG

CACATGCTGTGAGCTCGATGAGTAGGGCGGGACACGTGGTATCCTGTCTGAATATGGGGGGACCATCCTC

CAAGGCTAAATACTCCTGACTGACCGATAGTGAACCAGTACCGTGAGGGAAAGGCGAAAAGAACCCCGGC

GAGGGGAGTGAAAAAGAACCTGAAACCGTGTACGTACAAGCAGTGGGAGCACGCTTAGGCGTGTGACTGC

GTACCTTTTGTATAATGGGTCAGCGACTTATATTCTGTAGCAAGGTTAACCGAATAGGGGAGCCGAAGGG

AAACCGAGTCTTAACTGGGCGTTAAGTTGCAGGGTATAGACCCGAAACCCGGTGATCTAGCCATGGGCAG

GTTGAAGGTTGGGTAACACTAACTGGAGGACCGAACCGACTAATGTTGAAAAATTAGCGGATGACTTGTG

GCTGGGGGTGAAAGGCCAATCAAACCGGGAGATAGCTGGTTCTCCCCGAAAGCTATTTAGGTAGCGCCTC

GTGAATTCATCTCCGGGGGTAGAGCACTGTTTCGGCAAGGGGGTCATCCCGACTTACCAACCCGATGCAA

ACTGCGAATACCGGAGAATGTTATCACGGGAGACACACGGCGGGTGCTAACGTCCGTCGTGAAGAGGGAA

ACAACCCAGACCGCCAGCTAAGGTCCCAAAGTCATGGTTAAGTGGGAAACGATGTGGGAAGGCCCAGACA

GCCAGGATGTTGGCTTAGAAGCAGCCATCATTTAAAGAAAGCGTAATAGCTCACTGGTCGAGTCGGCCTG

CGCGGAAGATGTAACGGGGCTAAACCATGCACCGAAGCTGCGGCAGCGACACTATGTGTTGTTGGGTAGG

GGAGCGTTCTGTAAGCCTGTGAAGGTGTGCTGTGAGGCATGCTGGAGGTATCAGAAGTGCGAATGCTGAC

ATAAGTAACGATAAAGCGGGTGAAAAGCCCGCTCGCCGGAAGACCAAGGGTTCCTGTCCAACGTTAATCG

GGGCAGGGTGAGTCGACCCCTAAGGCGAGGCCGAAAGGCGTAGTCGATGGGAAACAGGTTAATATTCCTG

TACTTGGTGTTACTGCGAAGGGGGGACGGAGAAGGCTATGTTGGCCGGGCGACGGTTGTCCCGGTTTAAG

CGTGTAGGCTGGTTTTCCAGGCAAATCCGGAAAATCAAGGCTGAGGCGTGATGACGAGGCACTACGGTGC

TGAAGCAACAAATGCCCTGCTTCCAGGAAAAGCCTCTAAGCATCAGGTAACATCAAATCGTACCCCAAAC

CGACACAGGTGGTCAGGTAGAGAATACCAAGGCGCTTGAGAGAACTCGGGTGAAGGAACTAGGCAAAATG

GTGCCGTAACTTCGGGAGAAGGCACGCTGATATGTAGGTGAAGCGACTTGCTCGTGGAGCTGAAATCAGT

CGAAGATACCAGCTGGCTGCAACTGTTTATTAAAAACACAGCACTGTGCAAACACGAAAGTGGACGTATA

CGGTGTGACGCCTGCCCGGTGCCGGAAGGTTAATTGATGGGGTTAGCCGCAAGGCGAAGCTCTTGATCGA

AGCCCCGGTAAACGGCGGCCGTAACTATAACGGTCCTAAGGTAGCGAAATTCCTTGTCGGGTAAGTTCCG

ACCTGCACGAATGGCGTAATGATGGCCAGGCTGTCTCCACCCGAGACTCAGTGAAATTGAACTCGCTGTG

AAGATGCAGTGTACCCGCGGCAAGACGGAAAGACCCCGTGAACCTTTACTATAGCTTGACACTGAACATT

GAGCCTTGATGTGTAGGATAGGTGGGAGGCTTTGAAGTGTGGACGCCAGTCTGCATGGAGCCGACCTTGA

AATACCACCCTTTAATGTTTGATGTTCTAACGTTGACCCGTAATCCGGGTTGCGGACAGTGTCTGGTGGG

TAGTTTGACTGGGGCGGTCTCCTCCTAAAGAGTAACGGAGGAGCACGAAGGTTGGCTAATCCTGGTCGGA

CATCAGGAGGTTAGTGCAATGGCATAAGCCAGCTTGACTGCGAGCGTGACGGCGCGAGCAGGTGCGAAAG

CAGGTCATAGTGATCCGGTGGTTCTGAATGGAAGGGCCATCGCTCAACGGATAAAAGGTACTCCGGGGAT

AACAGGCTGATACCGCCCAAGAGTTCATATCGACGGCGGTGTTTGGCACCTCGATGTCGGCTCATCACAT

CCTGGGGCTGAAGTAGGTCCCAAGGGTATGGCTGTTCGCCATTTAAAGTGGTACGCGAGCTGGGTTTAGA

ACGTCGTGAGACAGTTCGGTCCCTATCTGCCGTGGGCGCTGGAGAACTGAGGGGGGCTGCTCCTAGTACG

AGAGGACCGGAGTGGACGCATCACTGGTGTTCGGGTTGTCATGCCAATGGCACTGCCCGGTAGCTAAATG

CGGAAGAGATAAGTGCTGAAAGCATCTAAGCACGAAACTTGCCCCGAGATGAGTTCTCCCTGACTCCTTG

AGAGTCCTGAAGGAACGTTGAAGACGACGACGTTGATAGGCCGGGTGTGTAAGCGCAGCGATGCGTTGAG

CTAACCGGTACTAATGAACCGTGAGGCTTAACCTT

**AAAAAAAAAAAA**

Modified-C-containing RNA strands in which the k-mers studied are underlined.

##### 5’-AAGCTAATACGACTCACTATAG

GAGTATAGGATTAGATAGAT<u>TGCTT</u>AGTTGAAGTATAGTAGATTAGA<u>GTCAT</u>AGAAGATGAGATTGAG<u>TGCAA</u>TTA

GAAGTTGATGTATAGATGA<u>ATCTG</u>TTAGATGGATAGTAATTAGTAGAGATTGAAG<u>ATCTT</u>GTAGATATGTTAGTA<u>T</u> <u>TCGA</u>TGATGAGGTGATATT<u>GTCGT</u>GGATATTAGATATATGGAGATGATAGTAGAGGATTGAAAAT**AAAAAAAAAA AA**

Modified-G-containing RNA strands in which the k-mers studied are underlined.

##### 5’-AAGCTAATACGACTCACTATAG

GACTATACCATTACATACAT<u>CTGCC</u>ACTTCACTCTACTACACTACA<u>ATGAA</u>TCCACATCACACTCAC<u>TAGAA</u>CTAC AACTCATCTACACATCA<u>ATGTT</u>CCACATCCATACTAACTACACACTCCAC<u>TTGTA</u>CTACATATCTCACTA<u>ATGCA</u>TC ATCACCTCATACT<u>TAGCC</u>ATATTACATCTACACATCATACTACACCACTCAAAAT**AAAAAAAAAAAA**

### Custom-modified NTP synthesis and characterization

The ribonucleosides m^2^G, m^5^U, and m^4^C_m_ used in the synthesis of the modified NTPs were obtained from commercial sources and used without further purification. The ribonucleoside (7.5 μmol) and 1,8-bis(dimethylamino)naphthalene (3.2 mg, 15 μmol) were suspended in trimethylphosphate (100 μL). The mixture was stirred at 4 °C for 5 min before phosphorus oxychloride (3 μL, 30 μmol) was pipetted into the mixture. The reaction was stirred for 2 h at 4 °C. A solution of tributylamine (2.5 μL) and tributylammonium pyrophosphate (5.5 mg, 10 μmol) in dry dimethylformamide (50 μL) was added and the mixture was stirred for another 2 h at 4 °C. The reaction was then quenched with triethylammonium bicarbonate solution (100 mM, 0.5 mL, pH 8) and stirred at room temperature for 30 min. The product was lyophilized, and the residue was carefully washed twice with ethyl acetate (400 μL). The white solid was dissolved in water and the product was purified by reversed-phase HPLC (A line: 50 mM triethylammonium bicarbonate pH 7; B line: acetonitrile; gradient: 0-15% MeCN over 30 min; flow rate: 1 mL/min; Abs = 260 nm) to afford the corresponding triphosphates. The purity of the triphosphates was determined by reversed-phase HPLC using the same method described above to be >90%. The presence of the triphosphate on the nucleoside was verified by ^31^P NMR (202 MHz, D_2_O): m^5^UTP δ -12.99 (m), -13.88 (m), -25.44 (m); m^2^GTP δ -12.61 (m), -13.45 (m), -25.34 (m); and m^4^C_m_TP δ -10.65 (m), -11.41 (m), -23.37 (m). The final yields were 17% for m^5^UTP, 15% for m^2^GTP, and 8% for m^4^C_m_TP.

### RNA library preparation

The purified RNA strands were first 3’-poly-A tailed with a commercial kit (Life Sciences) to allow the first step of library preparation to occur that ligates on a poly-T adaptor. The SQK-RNA002 kit provided by ONT was used without modification to the protocol. A noteworthy step in the library preparation method is the reverse transcription of the RNA by SuperScript III to form a RNA:DNA heteroduplex that facilitates longer read lengths to be obtained when conducting direct RNA nanopore sequencing. The prepared RNA libraries were loaded on either a MinION flow cell (∼10 ng) or a Flongle flow cell (∼5 ng) and sequenced using the standard settings in the MinKnow software. The ionic current vs. time trace data were stored in fast5 file formats that used the standard pass cutoff (Q > 7).

### Data analysis

The passed data were base called using Guppy (v 6.0.7). A newer version of Guppy exists and was not used in this work; however, in a previous report, we found Guppy v 6.3.2 gave similar miscalls for Ψ as v 6.0.7 and the impact of a newer base caller will likely not significantly change the results.^34^ The reads were aligned to the reference using minimap2 (command line for Tombo and ELIGOS2 analyses = “-ax map-ont -L”; Nanopore-Psu = “-ax splice -uf -k14”),^22^ sorted with SAM tools,^23^ and visualized with IGV.^24^ The references include the 16S and 23S rRNA sequences from the *rrnA* operon (see above in the IVT synthesis section). For the operon-specific Ψ analysis, the fastq reads were mapped to the *E. coli* strain K-12 substrain MG1655 genome (GenBank: U00096.2) as a reference. The operon expression levels were determined by the read density for the 16S rRNA sequences. The base-called data were quantified using either Nanopore-Psu^14^ or ELIGOS2^13^ following the procedures provided with these tools. The passed fast5 files were analyzed using Nanopolish,^25^ Nanocompore,^15^ and Tombo^26^ as described in the literature citations for these tools. All data were visualized using Origin or Excel.

### Associated Content Notes

A.M.F. and C.J.B. have a patent licensed to Electronic BioSciences and A.M.F. occasionally consuls on nucleic acid chemistry for Electronic BioSciences.

## Data Availability

The data and a complete supporting information will be available upon publication in a peer-reviewed journal.

## Acknowledgments

The National Institutes of Health provided financial support for this project (R01 GM093099 and R35 GM145237). We are grateful to Prof. Vahe Bandarian (University of Utah) for providing the *E. coli* knock-out strains from the Keio collection stored in his laboratory.

## References

(1) Jones, J. D.; Monroe, J.; Koutmou, K. S. A molecular-level perspective on the frequency, distribution, and consequences of messenger RNA modifications. Wiley Interdiscip. Rev. RNA 2020, e1586.

(2) Watson, Z. L.; Ward, F. R.; Méheust, R.; Ad, O.; Schepartz, A.; Banfield, J. F.; Cate, J. H. Structure of the bacterial ribosome at 2 Å resolution. eLife 2020, 9, e60482.

(3) Shi, H.; Wei, J.; He, C. Where, when, and how: Context-dependent functions of RNA methylation writers, readers, and erasers. Mol. Cell 2019, 74, 640–650.

(4) Thompson, M. G.; Sacco, M. T.; Horner, S. M. How RNA modifications regulate the antiviral response. Immunol. Rev. 2021, 304, 169–180.

(5) Su, D.; Chan, C. T.; Gu, C.; Lim, K. S.; Chionh, Y. H.; McBee, M. E.; Russell, B. S.; Babu, I. R.; Begley, T. J.; Dedon, P. C. Quantitative analysis of ribonucleoside modifications in tRNA by HPLC-coupled mass spectrometry. Nat. Protoc. 2014, 9, 828–841.

(6) Dai, Q.; Zhang, L.-S.; Sun, H.-L.; Pajdzik, K.; Yang, L.; Ye, C.; Ju, C.-W.; Liu, S.; Wang, Y.; Zheng, Z.; Zhang, L.; Harada, B. T.; Dou, X.; Irkliyenko, I.; Feng, X.; Zhang, W.; Pan, T.; He, C. Quantitative sequencing using BID-seq uncovers abundant pseudouridines in mammalian mRNA at base resolution. Nat. Biotechnol. 2022, doi.org/10.1038/s41587-41022-01505-w.

(7) Hawley, B. R.; Jaffrey, S. R. Transcriptome-wide mapping of m(6) A and m(6) Am at single-nucleotide resolution using miCLIP. Curr. Protoc. Mol. Biol. 2019, 126, e88.

(8) Khoddami, V.; Yerra, A.; Mosbruger, T. L.; Fleming, A. M.; Burrows, C. J.; Cairns, B. R. Transcriptome-wide profiling of multiple RNA modifications simultaneously at single-base resolution. Proc. Nat. Acad. Sci. U.S.A. 2019, 116, 6784–6789.

(9) Branton, D.; Deamer, D.: Nanopore Sequencing An Introduction; World Scientific Publishing Co. Pte. Ltd., 2019.

(10) Begik, O.; Lucas, M. C.; Pryszcz, L. P.; Ramirez, J. M.; Medina, R.; Milenkovic, I.; Cruciani, S.; Liu, H.; Vieira, H. G. S.; Sas-Chen, A.; Mattick, J. S.; Schwartz, S.; Novoa, E. M. Quantitative profiling of pseudouridylation dynamics in native RNAs with nanopore sequencing. Nat. Biotechnol. 2021, 39, 1278–1291.

(11) Smith, A. M.; Jain, M.; Mulroney, L.; Garalde, D. R.; Akeson, M. Reading canonical and modified nucleobases in 16S ribosomal RNA using nanopore native RNA sequencing. PloS One 2019, 14, e0216709.

(12) Fleming, A. M.; Mathewson, N. J.; Howpay Manage, S. A.; Burrows, C. J. Nanopore dwell time analysis permits sequencing and conformational assignment of pseudouridine in SARS-CoV-2. ACS Cent. Sci. 2021, 7, 1707–1717.

(13) Jenjaroenpun, P.; Wongsurawat, T.; Wadley, T. D.; Wassenaar, Trudy M.; Liu, J.; Dai, Q.; Wanchai, V.; Akel, N. S.; Jamshidi-Parsian, A.; Franco, A. T.; Boysen, G.; Jennings, M. L.; Ussery, D. W.; He, C.; Nookaew, I. Decoding the epitranscriptional landscape from native RNA sequences. Nucleic Acids Res. 2020, 49, e7.

(14) Huang, S.; Zhang, W.; Katanski, C. D.; Dersh, D.; Dai, Q.; Lolans, K.; Yewdell, J.; Eren, A. M.; Pan, T. Interferon inducible pseudouridine modification in human mRNA by quantitative nanopore profiling. Genome Biol. 2021, 22, 330.

(15) Leger, A.; Amaral, P. P.; Pandolfini, L.; Capitanchik, C.; Capraro, F.; Miano, V.; Migliori, V.; Toolan-Kerr, P.; Sideri, T.; Enright, A. J.; Tzelepis, K.; van Werven, F. J.; Luscombe, N. M.; Barbieri, I.; Ule, J.; Fitzgerald, T.; Birney, E.; Leonardi, T.; Kouzarides, T. RNA modifications detection by comparative nanopore direct RNA sequencing. Nat. Commun. 2021, 12, 7198.

(16) Tavakoli, S.; Nabizadeh, M.; Makhamreh, A.; Gamper, H.; McCormick, C. A.; Rezapour, N. K.; Hou, Y.-M.; Wanunu, M.; Rouhanifard, S. H. Semi-quantitative detection of pseudouridine modifications and type I/II hypermodifications in human mRNAs using direct long-read sequencing. Nat. Commun. 2023, 14, 334.

(17) Ramasamy, S.; Sahayasheela, V. J.; Sharma, S.; Yu, Z.; Hidaka, T.; Cai, L.; Thangavel, V.; Sugiyama, H.; Pandian, G. N. Chemical probe-based nanopore sequencing to selectively assess the RNA modifications. ACS Chem. Biol. 2022, 17, 2704–2709.

(18) Stephenson, W.; Razaghi, R.; Busan, S.; Weeks, K. M.; Timp, W.; Smibert, P. Direct detection of RNA modifications and structure using single-molecule nanopore sequencing. Cell Genom. 2022, 2, 100097.

(19) Thomas, N. K.; Poodari, V. C.; Jain, M.; Olsen, H. E.; Akeson, M.; Abu-Shumays, R. L. Direct nanopore sequencing of individual full length tRNA strands. ACS Nano 2021, 15, 16642–16653.

(20) Popova, A. M.; Williamson, J. R. Quantitative analysis of rRNA modifications using stable isotope labeling and mass spectrometry. J. Am. Chem. Soc. 2014, 136, 2058–2069.

(21) Kurylo, C. M.; Parks, M. M.; Juette, M. F.; Zinshteyn, B.; Altman, R. B.; Thibado, J. K.; Vincent, C. T.; Blanchard, S. C. Endogenous rRNA sequence variation can regulate stress response gene expression and phenotype. Cell Rep. 2018, 25, 236–248.

(22) Li, H. Minimap2: pairwise alignment for nucleotide sequences. Bioinformatics 2018, 34, 3094–3100.

(23) Li, H.; Handsaker, B.; Wysoker, A.; Fennell, T.; Ruan, J.; Homer, N.; Marth, G.; Abecasis, G.; Durbin, R. The sequence alignment/map format and SAMtools. Bioinformatics 2009, 25, 2078–2079.

(24) Robinson, J. T.; Thorvaldsdóttir, H.; Winckler, W.; Guttman, M.; Lander, E. S.; Getz, G.; Mesirov, J. P. Integrative genomics viewer. Nat. Biotechnol. 2011, 29, 24–26.

(25) Loman, N. J.; Quick, J.; Simpson, J. T. A complete bacterial genome assembled de novo using only nanopore sequencing data. Nat. Meth. 2015, 12, 733–735.

(26) Stoiber, M.; Quick, J.; Egan, R.; Eun Lee, J.; Celniker, S.; Neely, R. K.; Loman, N.; Pennacchio, L. A.; Brown, J. De novo identification of DNA modifications enabled by genome-guided nanopore signal processing. bioRxiv 2017, 094672.

(27) Baba, T.; Ara, T.; Hasegawa, M.; Takai, Y.; Okumura, Y.; Baba, M.; Datsenko, K.; Tomita, M.; Wanner, B.; Mori, H. Construction of Escherichia coli K-12 in-frame, single-gene knockout mutants: the Keio collection. Mol. Syst. Biol. 2006, 2, 2006.0008.

(28) Sergeeva, O. V.; Bogdanov, A. A.; Sergiev, P. V. What do we know about ribosomal RNA methylation in Escherichia coli? Biochimie 2015, 117, 110–118.

(29) Kimura, S.; Sakai, Y.; Ishiguro, K.; Suzuki, T. Biogenesis and iron-dependency of ribosomal RNA hydroxylation. Nucleic Acids Res. 2017, 45, 12974–12986.

(30) Bakin, A.; Kowalak, J. A.; McCloskey, J. A.; Ofengand, J. The single pseudouridine residue in Escherichia coli 16S RNA is located at position 516. Nucleic Acids Res. 1994, 22, 3681–3684.

(31) Conrad, J.; Sun, D.; Englund, N.; Ofengand, J. The rluC gene of Escherichia coli codes for a pseudouridine synthase that is solely responsible for synthesis of pseudouridine at positions 955, 2504, and 2580 in 23 S ribosomal RNA. The Journal of biological chemistry 1998, 273, 18562–18566.

(32) Del Campo, M.; Kaya, Y.; Ofengand, J. Identification and site of action of the remaining four putative pseudouridine synthases in Escherichia coli. RNA 2001, 7, 1603–1615.

(33) Lesnyak, D. V.; Sergiev, P. V.; Bogdanov, A. A.; Dontsova, O. A. Identification of Escherichia coli m^2^G methyltransferases: I. the ycbY gene encodes a methyltransferase specific for G2445 of the 23 S rRNA. J. Mol. Biol. 2006, 364, 20–25.

(34) Fleming, A. M.; Burrows, C. J. Nanopore sequencing for N1-methylpseudouridine in RNA reveals sequence-dependent discrimination of the modified nucleotide triphosphate during transcription. Nucleic Acids Res. 2023, 51, 1914–1926.

(35) René, O.; Alix, J. H. Late steps of ribosome assembly in E. coli are sensitive to a severe heat stress but are assisted by the HSP70 chaperone machine. Nucleic Acids Res. 2011, 39, 1855–1867.

(36) Cheng-Guang, H.; Gualerzi, C. O. The ribosome as a switchboard for bacterial stress response. Front. Microbiol. 2020, 11, 619038.

(37) Liu, H.; Begik, O.; Lucas, M. C.; Ramirez, J. M.; Mason, C. E.; Wiener, D.; Schwartz, S.; Mattick, J. S.; Smith, M. A.; Novoa, E. M. Accurate detection of m6A RNA modifications in native RNA sequences. Nat. Commun. 2019, 10, 4079.

(38) Craig, J. M.; Laszlo, A. H.; Brinkerhoff, H.; Derrington, I. M.; Noakes, M. T.; Nova, I. C.; Tickman, B. I.; Doering, K.; de Leeuw, N. F.; Gundlach, J. H. Revealing dynamics of helicase translocation on single-stranded DNA using high-resolution nanopore tweezers. Proc. Natl. Acad. Sci. U. S. A. 2017, 114, 11932–11937.

(39) Nookaew, I.; Jenjaroenpun, P.; Du, H.; Wang, P.; Wu, J.; Wongsurawat, T.; Moon, S. H.; Huang, E.; Wang, Y.; Boysen, G. Detection and discrimination of DNA adducts differing in size, regiochemistry, and functional group by nanopore sequencing. Chem. Res. Toxicol. 2020, 33, 2944–2952.

(40) Heron, A.; Clarke, J.; Moysey, R.; Wallace, J.; Bruce, M.; Jayasinghe, L.; Caprotti, D.; Soeroes, S.; McNeill, L.; Alves, D.; Bowen, R.; Milton, J. Modified Helicases US2015/0191709A1, Jul. 9, 2015.

(41) Büttner, K.; Nehring, S.; Hopfner, K. P. Structural basis for DNA duplex separation by a superfamily-2 helicase. Nat. Struct. Mol. Biol. 2007, 14, 647–652.

(42) Abou Assi, H.; Rangadurai, A. K.; Shi, H.; Liu, B.; Clay, M. C.; Erharter, K.; Kreutz, C.; Holley, C. L.; Al-Hashimi, Hashim M. 2′-O-Methylation can increase the abundance and lifetime of alternative RNA conformational states. Nucleic Acids Res. 2020, 48, 12365–12379.

(43) Kimura, S.; Suzuki, T. Fine-tuning of the ribosomal decoding center by conserved methyl-modifications in the Escherichia coli 16S rRNA. Nucleic Acids Res. 2010, 38, 1341–1352.

(44) Fasnacht, M.; Gallo, S.; Sharma, P.; Himmelstoß, M.; Limbach, Patrick A.; Willi, J.; Polacek, N. Dynamic 23S rRNA modification ho5C2501 benefits Escherichia coli under oxidative stress. Nucleic Acids Res. 2021, 50, 473–489.

(45) Rivière, J.; Klarskov, K.; Wagner, J. R. Oxidation of 5-hydroxypyrimidine nucleosides to 5-hydroxyhydantoin and its alpha-hydroxy-ketone isomer. Chem. Res. Toxicol. 2005, 18, 1332–1338.

(46) O’Connor, M.; Leppik, M.; Remme, J. Pseudouridine-free Escherichia coli ribosomes. J. Bacteriol. 2018, 200, e00540.

(47) Schmidt, A.; Kochanowski, K.; Vedelaar, S.; Ahrné, E.; Volkmer, B.; Callipo, L.; Knoops, K.; Bauer, M.; Aebersold, R.; Heinemann, M. The quantitative and condition-dependent Escherichia coli proteome. Nat. Biotechnol. 2016, 34, 104–110.

(48) Huber, S. M.; Leonardi, A.; Dedon, P. C.; Begley, T. J. The versatile roles of the tRNA epitranscriptome during cellular responses to toxic exposures and environmental stress. Toxics 2019, 7.

